# An evolutionarily conserved pacemaker role for HCN ion channels in smooth muscle

**DOI:** 10.1101/2022.08.14.503722

**Authors:** Lei Yang, Rodolfo J. Ricart Arbona, Carl Smith, Kelly M. Banks, V. Kaye Thomas, Lawrence Palmer, Todd Evans, Romulo Hurtado

## Abstract

Although HCN ion channels are well established to underlie cardiac pacemaker activity, their role in smooth muscle organs remains controversial. HCN expressing cells are localized to renal pelvic smooth muscle (RPSM) pacemaker tissues of the murine upper urinary tract and HCN channel conductance is required for peristalsis. To date, however, the *I*_h_ pacemaker current conducted by HCN channels has never been detected in these cells, raising questions on the identity of RPSM pacemakers. Indeed, the RPSM pacemaker mechanisms of the unique multicalyceal upper urinary tract exhibited by humans remains unknown. Here, we developed immunopanning purification protocols and demonstrate that 96% of isolated HCN+ cells exhibit *I*_h_. Single molecule STORM to whole-tissue imaging showed HCN+ cells express single HCN channels on their plasma membrane and integrate into the muscular syncytium. By contrast, PDGFR-α+ cells exhibiting the morphology of ICC gut pacemakers were shown to be vascular mural cells. Translational studies in the homologous human and porcine multicalyceal upper urinary tracts showed that contractions and pacemaker depolarizations originate in proximal calyceal RPSM. Critically, HCN+ cells were shown to integrate into calyceal RPSM pacemaker tissues, and HCN channel block abolished electrical pacemaker activity and peristalsis of the multicalyceal upper urinary tract. Cumulatively, these studies demonstrate that HCN ion channels play a broad, evolutionarily conserved pacemaker role in both cardiac and smooth muscle organs and have implications for channelopathies as putative etiologies of smooth muscle disorders.

## INTRODUCTION

Synchronized contractile activity of involuntary muscles is critical for animal and human physiology. In the heart, atrial-to-ventricular contractions are essential for efficient transport of blood throughout the body. In tubular smooth muscles, such as in the gastrointestinal (GI) and upper urinary tract (UUT), proximal-to-distal contractions are necessary to propel luminal contents forward. The coordinated, myogenic contractions of these autonomic muscles are triggered by intrinsic pacemakers ^1-3^. The anatomic position of pacemakers is critical, as it sets the origin of propagating contractile waves. Indeed, ectopic pacemaker activity results in debilitating muscle dysfunction, such as cardiac and smooth muscle arrhythmias ^4-6^.

Hyperpolarization-activated cation (HCN) channels are conserved and underlie cardiac pacemaker activity in all animal species ^7^. Unlike the majority of voltage-gated ion channels, HCN channel family members (HCN1-4) conduct an inward cation current that is activated by membrane hyperpolarization (*I*_h_) to potentials more negative than -45 mV. It is this unusual biophysical property of HCN channels that enable *I*_h_ to elicit rhythmic pacemaker depolarizations. *I*_h_ is inward in the activation range of HCN channels, which is near or below the diastolic potential of cells. Thus, after an action potential, *I*_h_ depolarizes the membrane back toward threshold for triggering another action potential, thereby establishing self-sustaining repetitive firing ^7^. The mammalian sinoatrial node (SAN), the pacemaker of the heart, is localized along the terminal crest of the right atrium and is marked by the expression of HCN2 and HCN4 ^8-10^. Loss-of-function and gain-of-function studies have firmly established that *I*_h_ underlies SAN pacemaker activity that drives a coordinated heartbeat. Pharmacological inhibition with HCN blockers, including Cs^+^ and ZD7288, as well as HCN2/4 genetic ablation result in perturbed pacemaker excitation and abnormal heart contractions ^11-14^. In humans, HCN4 channelopathies are associated with lethal cardiac arrhythmias ^4, 15^. Finally, pacemaker activity can be induced in electrically quiescent cells transfected with HCN channels, and bioengineered HCN-expressing cells integrate into and drive beating of donor hearts ^16-18^.

There is a generally held assumption that cardiac and smooth muscle pacemaker mechanisms must be distinct due to quantitative differences in their contractile rhythms, such as a smooth muscle contractile rate that is an order of magnitude slower. Recent studies, however, provide evidence against this dogma, suggesting a role for HCN channels in regulating smooth muscle systems ^7, 19^. Pacemaker tissues localize to the kidney proximal end of the UUT, termed the renal pelvis. HCN channels are expressed in renal pelvic smooth muscle (RPSM) pacemaker tissues of the UUT in numerous mammalian species, including humans ^20-25^. Live imaging of electrical and contractile excitation through the intact murine UUT has shown that HCN channel conductance is required for the generation of RPSM pacemaker activity and coordinated UUT peristalsis that propels waste away from the kidney ^20^. Inhibition of HCN channel conductance also abolishes spontaneous bursting activity of individual HCN+ cells localized to RPSM pacemaker tissues ^26^. HCN^+^ RPSM cells are electrically coupled to the smooth muscle and co-express T-type calcium channels that mediate HCN-dependent pacemaker activity and modulate the rate of UUT peristalsis ^23^. Moreover, perturbed UUT peristalsis and UUT dysfunction is exhibited by mutant mice lacking HCN^+^ cells ^21^.

Despite these findings, the role of HCN+ pacemakers in smooth muscle remains controversial. To date, the HCN channel hyperpolarization-activated (*I*_h_) current that drives pacemaker activity has yet to be detected in RPSM cells. This has raised doubt whether HCN+ cells are bona fide RPSM pacemakers ^19, 27^. Furthermore, recent studies in the UUT have identified Platelet Derived Growth Factor Receptor-α (PDGFR-α)+ cells exhibiting a morphology similar to ICC gut pacemakers ^3^, and based on these morphological similarities these cells were considered possible UUT pacemakers ^28^. However, PDGFR-α^+^ cells constituted a dynamic interconnected network of cells that spanned from the proximal UUT pacemaker tissue down to distal muscle segments, and they penetrated into all UUT tissue layers, including the urothelial, smooth muscle and adventitial layers. Moreover, PDGFR-α^+^ cells also formed an interconnected network of cells surrounding renal tubules in kidney tissue, thus raising questions regarding the identity of these cells. By contrast, these same studies demonstrated that clusters of HCN3+ cells are restricted to RPSM pacemaker tissues. Thus, the identity RPSM pacemakers remains to be resolved.

Interpreting the clinical significance of the aforementioned studies also comes with the additional caveat that higher order mammals such as swine and humans exhibit a multicalyceal UUT with a unique anatomy and physiology ^29-31^. In these species, rhythmic contractile and electrical excitation has been shown to emanate from the proximal calyceal RPSM ^32-34^. To date however, the mechanism underlying pacemaker activity of RPSM calyx pacemaker tissue remains unknown.

In this study, we perform comprehensive imaging studies from the single molecule to the whole tissue level to characterize putative pacemaker cells in both the unicalyceal and multicalyceal UUTs. Methods are established to purify RPSM pacemakers and characterize their electrophysiological properties. Finally, a model system is established that enables the continuous analysis of electrical and contractile waves through the intact multicalyceal UUT. Our studies demonstrate that HCN+ cells exhibit stereotypic electrophysiological properties of pacemakers, and that they make up part of the muscular syncytium of RPSM pacemaker tissues in the unicalyceal and multicalyceal UUT. By contrast, we show that PDGFR-α cells are vascular mural cells. Moreover, functional studies showed that HCN channel conductance underlies pacemaker activity that triggers coordinated peristalsis of the multicalyceal UUT.

## RESULTS

### Pacemaker tissues of the heart and UUT exhibit an anatomic architecture consisting of an HCN+ cell tissue layer integrated into the musculature

Understanding the cellular architecture of an organ provides critical insight into the function of different cell types. To this end, we performed whole tissue imaging of the UUT to better understand the roles of ICC and HCN+ cells. Indeed, given the parity in HCN channel expression, we hypothesized that the heart and UUT may exhibit analogous pacemaker tissue architectures. Imaging was first performed to validate the known anatomy of the heart. Whole murine atria were stained for cardiac Troponin (green) to mark the myocardium and HCN4 (red) to mark pacemakers. Heart anatomy was strikingly revealed, including the intricate interconnections and contours of the cardiac muscle fibers, and the localization of the SAN along the terminal crest of the right atrium (Fig. 1A). High magnification analyses detailed the adjacent localization of HCN+ pacemakers and myocardium in the SAN (Fig. 1A, inset and supplemental Movie 1).

**Figure 1.**
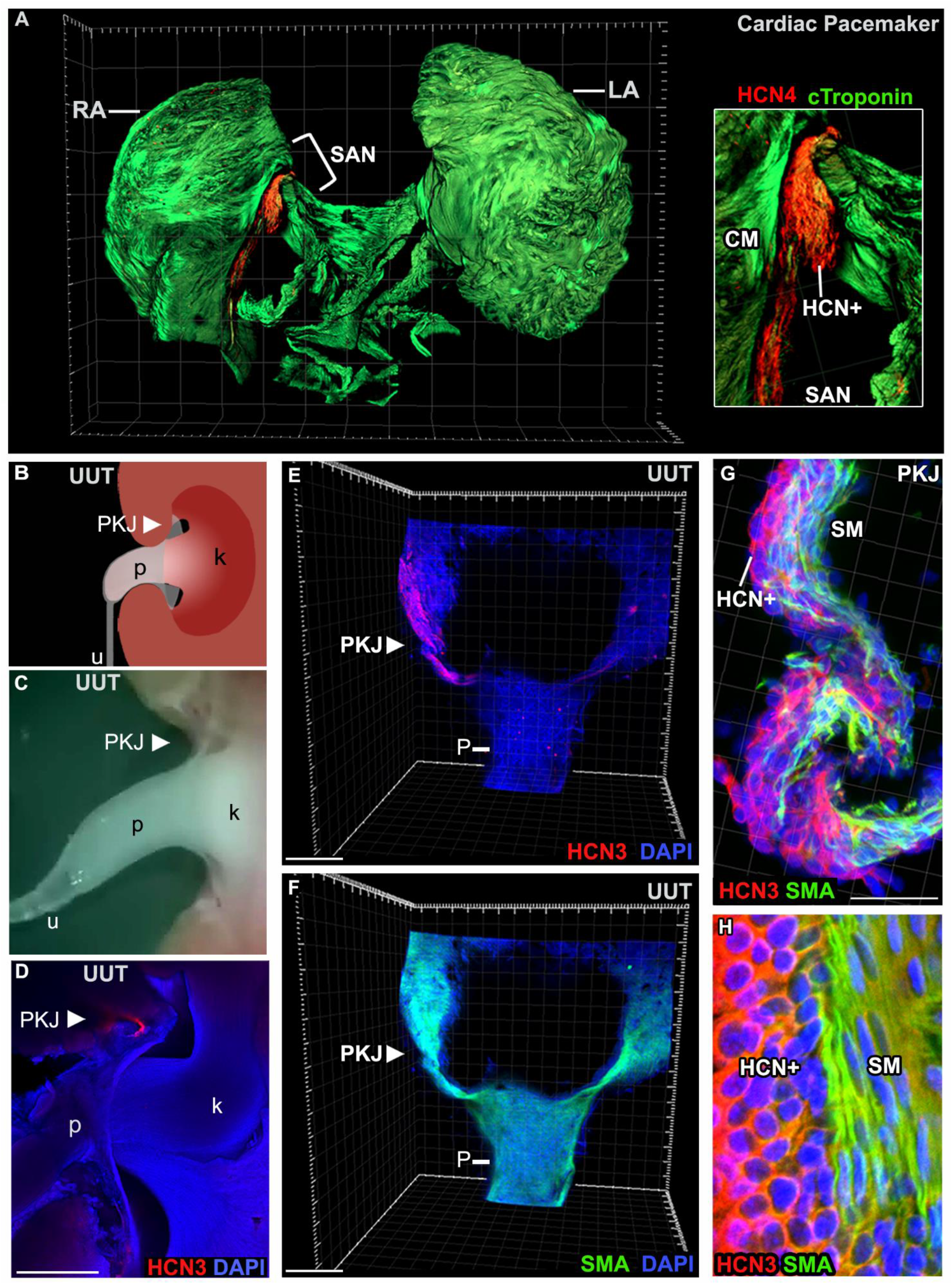
Cardiac and RPSM pacemaker tissues contain an analogous tissue architecture consisting of an HCN+ cell layer that makes up part of the musculature. **A**) Confocal imaging of whole adult murine atria immunolabeled for c-Troponin (green) to mark the myocardium and HCN4 to mark the SAN pacemaker (Inset, high magnification of the SAN). See supplemental movie 1. **B)** Schematic of the murine UUT, which is a continuous muscular organ with a flared proximal end termed the renal pelvis (p) and a more narrowed distal end, termed the ureter (u). RPSM pacemaker tissue localizes to the site where the pelvis joins the kidney (k), termed the pelvis kidney junction (PKJ). **C)** Representative still image of supplemental movie 2 showing live peristaltic contractions of the murine UUT. **D)** Cryosection of murine UUT immunostained for HCN3 (red) and DAPI. **E-F)** Imaging of whole proximal UUTs immunolabeled for HCN3 (E, red) and SMA (F, green). Low magnification analyses (E, F) revealed HCN+ cell tissue layer localized to the PKJ pacemaker. High magnification analyses showed that the HCN+ tissue layer was integrated into UUT musculature (G; see supplementary movie 3), with HCN+ cells and smooth muscle cells forming adjacent tissue layers (H).

The UUT is a continuous muscular organ with a flared proximal end, termed the renal pelvis, and a more narrowed distal end termed the ureter (Fig. 1B, schematic; 1C, still image of the murine UUT). In the UUT, pacemaker tissues localize to the site where the renal pelvis meets the kidney, termed the pelvis-kidney junction (PKJ). PKJ pacemaker activity triggers rhythmic, peristaltic contractions that emanate from the PKJ and propagate in a coordinated fashion distally down the UUT (supplemental Movie 2). Numerous groups have used standard IF of thin UUT sections to show that HCN3+ cells localize to the PKJ (Fig. 1D) ^20-23, 25, 26, 28^. However, these studies lack information about the gross anatomical architecture of the PKJ pacemaker tissue. To this end, we performed imaging of the intact UUT (Fig. 1E, F). Whole UUTs were stained for HCN3, SMA and DAPI, and then imaged by confocal microscopy. HCN+ cells localized to the PKJ (Fig. 1E, HCN3, red) of the UUT (Fig. 1F, SMA, green). Critically, high magnification analyses showed that HCN+ cells are integrated within the RPSM of the UUT (Fig. 1G, and supplemental Movie 3). Similar to the heart, HCN+ cells and smooth muscle cells formed adjacent tissue layers (Fig. 1H). Thus, we provide the first documentation of the gross, intact tissue architecture of the PKJ pacemaker and establish that HCN+ cells make up part of the RPSM musculature.

### PDGFR-α cells of the UUT are vascular associated mural cells

PDGFR-α+ cells were recently identified in the UUT and proposed to be pacemakers ^28^. However, these PDGFR-α+ cells formed an interconnected network of cells in both the kidney and UUT, and thus we hypothesized that they could be vascular associated mural cells. Indeed, PDGFR-α+ cells have already been established in the kidney to be vascular mural cells ^35^. To test our hypothesis, we first imaged the vasculature (endomucin, green) and PDGFR-α+ cells (red) of the kidney (supplemental Fig. 1A, white brackets in top panels denote areas imaged at high magnification in bottom panels). The known architecture of the peritubular vasculature was readily evident, with vessels surrounding the renal tubules (rt). Consistent with published studies, our imaging confirmed that PDGFR-α+ cells are renal vascular mural cells associated with the peritubular vasculature of the kidney.

Having confirmed the known cell identity of PDGFR-α+ cells in the kidney, we next determined whether PDGFR-α+ cells of the UUT are also vascular-associated mural cells. Intact UUTs were immunostained for the vasculature (endomucin, green) and PDGFR-α (red). As illustrated in three independent representative samples (Fig. 2), we discovered that PDGFR-α + cells of the UUT are indeed vascular mural cells. PDGFR-α cells overlaid the vascular tree of the UUT that spanned from the proximal PKJ pacemaker tissue (Fig.2A-H), down the RPSM to the distal UUT where the pelvis joins the ureter, or the UPJ (Fig. 2I-K). Consistent with the known spatial distribution of mural cells, PDGFR-α+ cells localized to large caliber vessels, whereas subsets of small caliber vessels lacked mural cell coverage (arrows, Fig. 2F-H, and L-N). To thoroughly examine the identity of PDGFR-α cells throughout the UUT, we assayed distal muscle segments of additional independent UUTs (Suppl. Fig. 1B). PDGFR-α+ cells were again found to cover the vascular bed of the renal pelvis and ureter at the UPJ. Taken together, our imaging studies establish that HCN+ cells of the UUT make up part of the UUT musculature, whereas PDGFR-α cells are vascular mural cells.

**Figure 2.**
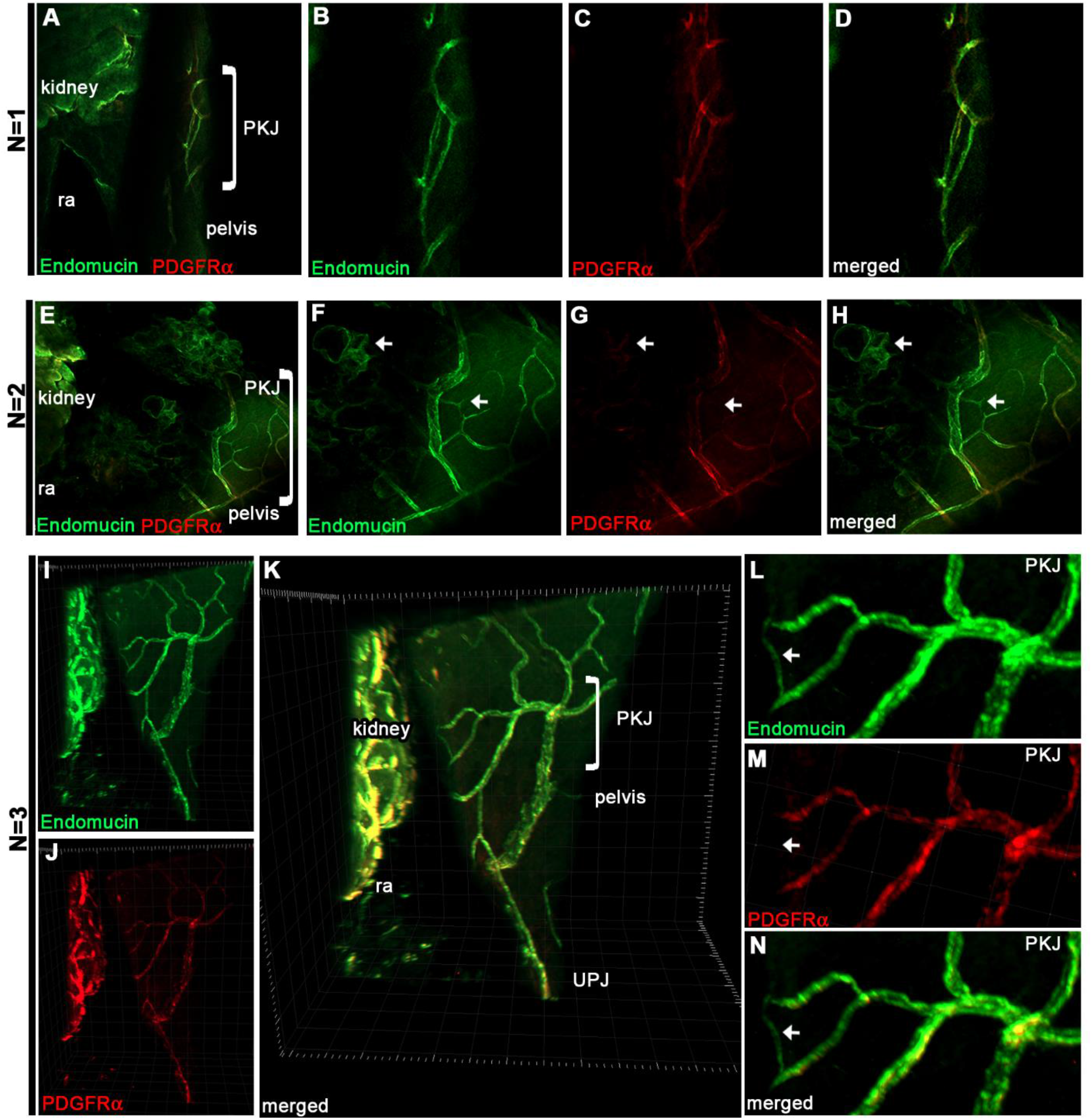
PDGFR-α+ cells of the UUT are vascular associated mural cells. **A-N)** The cell identity of PDGFR-α+ cells was assayed in the intact UUT. Three independent representative samples are shown (N=1, A-D; N=2, E-H; N=3, I-N). UUTs were immunolabeled for the vasculature (vascular endothelium, endomucin, green) and PDGFR-α (red). PDGFR-α+ cells proved to be vascular associated mural cells, overlaying the vascular tree of the UUT. Arrows point to small caliber vessels lacking PDGFR-α+ mural cell coverage (F-H, and L-N). Brackets in A, E, and K mark PKJ tissue that was assayed at higher magnification (B-D; F-H; and L-N). pelvis-kidney junction, PKJ; ureter pelvis junction, UPJ.

### Validation of HCN+ cell identity

Our imaging studies of the PKJ tissue architecture are consistent with a role for HCN+ cells as pacemakers. Thus, we performed further studies to validate the pacemaker function of HCN+ cells. Tissue separation studies demonstrated that peristalsis persisted in isolated proximal UUT tissues (see supplemental Movie 4). Isolated PKJ tissues that were dissected out also exhibited rhythmic myogenic contractions (see supplemental Movie 5). In contrast, proximal UUT segments imaged immediately after being disconnected from the PKJ lacked rhythmic contractility (see supplemental Movie 6). Critically, inhibition with ZD7288, the most widely used and characterized HCN channel blocker ^10-13, 36, 37^, abolished myogenic contractions of PKJ tissues segments (supplemental Movie 7, before inhibition; supplemental Movie 8, post inhibition), in agreement with published studies ^21, 23, 26^. Thus, HCN channels are required for initiating peristalsis in PKJ tissues.

To corroborate that whole tissue imaging of the PKJ is indeed representative of ion channel expression in HCN+ cells, super resolution imaging was performed on isolated PKJ tissue and primary cells freshly dissociated from the PKJ (Fig. 3). Isolated PKJ tissues were cryosectioned, immunostained for HCN3 and analyzed by total internal reflection (TIRF) microscopy in combination with STORM super resolution analyses. TIRF microscopy of HCN3 staining imaged the cell membrane boundary at the interface between HCN3+ cells (Fig. 3A, HCN3; inset DAPI). STORM analyses of TIRF microscopy provided sub-diffraction limit resolution of HCN3 staining (Fig.3B) and facilitated the visualization of clusters and single HCN3 channels expressed by PKJ cells (Fig. 3C, D). Figures 3E and F show phase contrast images of typical plated primary cells freshly prepared from PKJ tissue. IF analyses of PKJ primary cells revealed that HCN3+ cells of the PKJ exhibit a morphology consistent with small, compact round cell types observed by phase contrast microscopy (Fig. 3G). TIRF and STORM imaging of compact, rounded primary PKJ cells was performed to determine the cellular localization of HCN channels in these cells. TIRF illumination to depths of approximately 150nm imaged channel expression localized to the plasma membrane (Fig. 3H), and STORM revealed clusters and single HCN3 channels on the cell surface of compact PKJ cells (Fig. 3I and J). Thus, HCN channels are expressed on the plasma membrane of PKJ cells.

**Figure 3.**
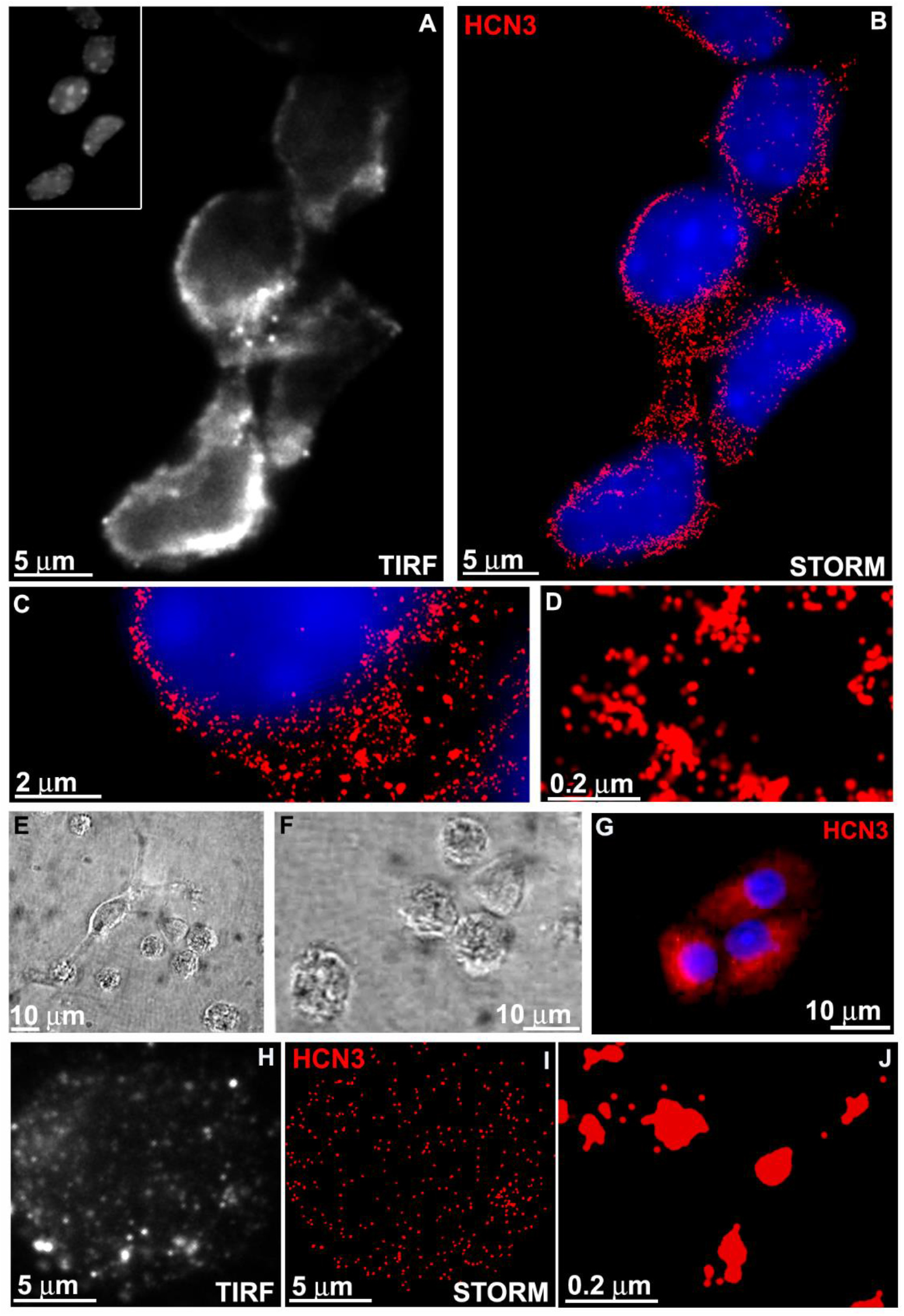
Super resolution imaging reveals HCN ion channels on the plasma membrane of PKJ cells with a compact, round morphology. **A-D)** Super resolution imaging of PKJ tissues immunolabeled for HCN3. (A) TIRF images of PKJ tissue immunolabeled for HCN3 (inset, DAPI). (B-D) STORM analyses of TIRF images (HCN3, red; blue, DAPI). **E, F)** Low (E) and high (F) magnification phase contrast images of primary cells freshly dissociated from the PKJ. **G)** IF of primary cells isolated from the PKJ (HCN3, red; DAPI, blue). HCN3+ cells exhibited a morphology consistent with small, compact round cell types observed by phase microscopy. **H-J)** Super resolution imaging of compact, rounded primary cells freshly isolated from the PKJ and immunolabeled for HCN3. (H) TIRF images of primary cells immunolabeled for HCN3. (I and J) STORM analyses of TIRF images (HCN3, red).

It has yet to be shown that PKJ cells exhibit the *I*_h_ pacemaker current conducted by HCN channels, which has raised concerns whether HCN+ cells are functional pacemakers ^19, 27, 28^. To this end, freshly isolated PKJ cells with the compact morphology of HCN3^+^ cells were examined using whole-cell patch clamp to examine their electrophysiological properties (Fig. 4). Since *I*_h_ is insensitive to Ba^2+^, we added 0.5mM Ba^2+^ to the bathing medium to block currents through K^+^ channels ^7-9^. To assay for *I*_h_, we used a holding potential of - 30 mV and applied hyperpolarizing voltage steps from -40 mV to -150 mV for at least 5 seconds. These cells expressed archetypal *I*_h_ currents with a characteristic slowly activating inward current that increased in amplitude with progressively negative voltages (Fig. 4A). Fig. 4B shows the I-V relationship of the hyperpolarization-activated current in these cells.

**Figure 4.**
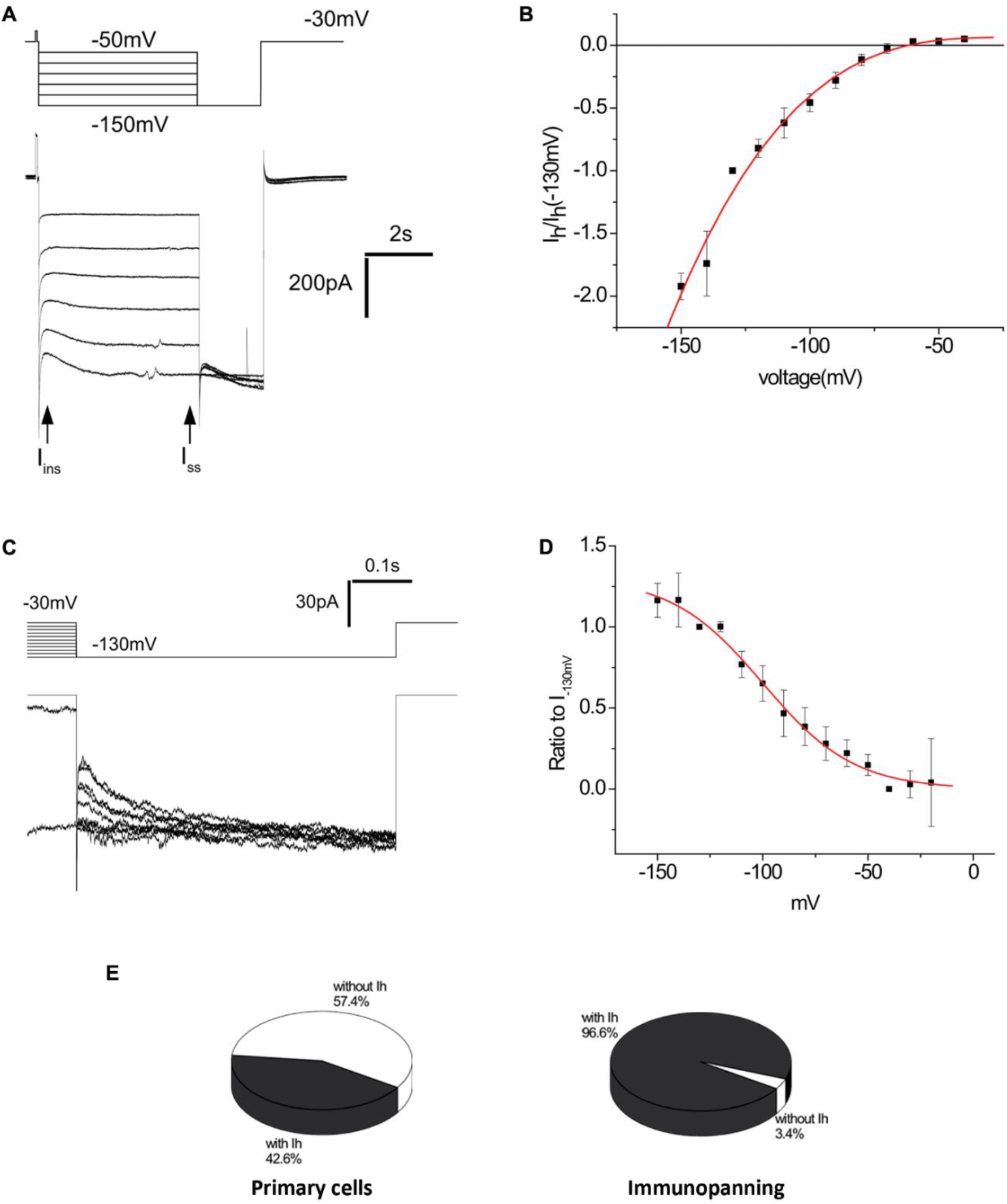
PKJ cells exhibit *I*_h_ current. **A-D)** Freshly isolated primary cells of the PKJ with the compact rounded morphology of HCN3+ cells were used for whole-cell voltage-clamp recording. (A) Typical current traces in response to hyperpolarization of the membrane potential from -50 mV. (B) voltage-dependence of *I*_h_, measured as the difference between the current at the end of the voltage step and at the beginning (after decay of the capacitance transient), normalized to the value at -130 mV. Data represent means ± SEM for 8 cells. (C) Voltage-dependent activation of *I*_h_ in PKJ cells. The voltage was clamped to values between -30 and -150 mV for 5 or 10 sec, and then held at -130 mV for 500 msec. (D) Data are plotted according to eq. 1 and normalized to values at -130 mV, V1/2 = -100 ± 3 mV and k = 22 ± 2. (E) Percentage of cells with measurable *I*_h_ in unselected primary cell populations (left diagram) and in primary cells purified by immunopanning with an anti-HCN3 antibody (right diagram).

A typical recording (Fig. 4C) and plot of the current amplitude at a fixed voltage (−130 mV), as a function of the preceding conditioning voltage (Fig. 4D), illustrates *I*_h_ activation in compact PKJ cells. Activation of *I*_h_ during the conditioning pulse increases the current at the beginning of the test pulse. *I*_h_ activated predominantly between -50 mV and -140 mV, reaching half-maximum at V_0.5_ = -100 ±3 mV. The time constant for activation at -130 mV was 440 ± 94 msec. These numbers are consistent with studies of HCN3 transfected into HEK293 cells, where V_0.5_ of *I*_h_ was -100 ±0.77 mV and the time constant was 470-570 msec ^38, 39^. Thus, the activation properties of *I*_h_ recorded from PKJ cells are consistent with the properties of heterologously expressed HCN channels. The mean current density at V = -130 mV was -5.5 ± 1.5 pA/pF.

*I*_h_ current was observed in 43% (23/54) of freshly isolated primary PKJ cells exhibiting a compact round morphology (Fig. 4E, primary cells), indicating heterogeneity of PKJ cells. To selectively isolate and analyze HCN+ cells of the PKJ, we developed an immunopanning protocol using an antibody to the extracellular domain of HCN3. Cells freshly isolated from the PKJ were transiently incubated on coverslips coated with anti-HCN3 antibody, and bound cells assayed by whole-cell patch clamp. In the immunopanned population we detected *I*_h_ in 96% (28/29) of cells, a dramatic increase over the unselected cells (Fig. 4E, immunopanning), thus confirming the correspondence between PKJ cells expressing HCN3 and those that exhibit *I*_h_.

The pharmacological properties of *I*_h_ in PKJ cells were analyzed using Cs^+^ and ZD7288, both well-characterized HCN channel blockers (Fig. 5) ^10, 11^. Fig. 5A shows representative currents from a cell before and after 13 min incubation with ZD7288, which decreased the amplitude of *I*_h_ by > 80% (Fig. 5B). Since ZD7288 is not amenable to washout, we performed additional experiments to preclude rundown effects. Specifically, purified compact PKJ cells were incubated with or without ZD7288 for at least 20 minutes prior to whole-cell patch clamp. Pre-incubation with ZD7288 also reduced *I*_h_ by >80% (Fig. 5C), confirming the inhibition of HCN channel conductance while avoiding channel rundown artifact. Representative traces from experiments using Cs^+^, a reversible HCN inhibitor, show > 95% inhibition of *I*_h_, which was partially reversed upon washout (Fig. 5D). Thus, these drug studies demonstrate that *I*_h_ expressed by PKJ cells exhibit the known pharmacological profiles of *I*_h_.

**Figure 5.**
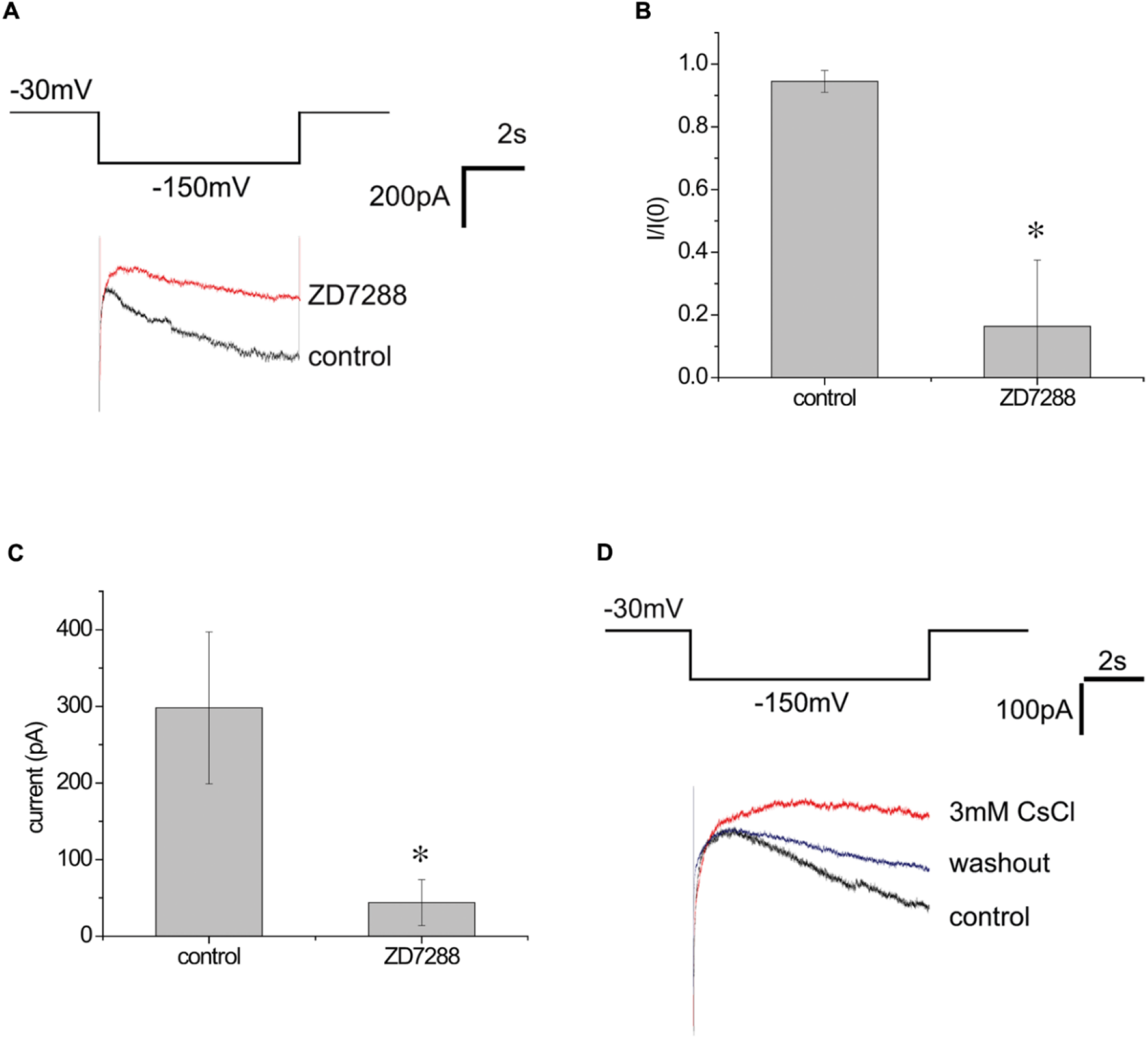
*I*_h_ current exhibited by PKJ cells is inhibited by HCN channel blockers. **A-C)** *I*_h_ expressed by cells of the PKJ is inhibited by the HCN channel antagonist ZD7288. (A) Representative trace of *I*_h_ before and 13 min after application of 100µM ZD7288. (B) Mean fractional change in *I*_h_ at V = -150 mV after inhibition with ZD7288 (mean ± SEM for n = 3 cells, *p= 0.05). (C) Mean values of *I*_h_ in immunopanned cells pre-incubated with or without 100µM ZD7288. (mean ± SEM n = 10 cells, *p= 0.05). (D) Block of *I*_h_ by 3 mM Cs^+^, which was partially reversible upon washout.

Cumulatively, our imaging, functional, and electrophysiological analyses demonstrate that HCN3+ PKJ cells are part of the muscular syncytium, that HCN channel conductance is required for smooth muscle peristalsis, that HCN channels are expressed on the plasma membrane of PKJ cells, and that PKJ cells exhibit the *I*_h_ pacemaker current typical of HCN channels. Taken together, these data provide strong evidence establishing HCN+ cells as the RPSM pacemakers.

### The HCN+ cells play a conserved pacemaker role in the multicalyceal UUT of higher order mammals

Higher-order mammalian species, including swine and humans, exhibit an UUT with a distinct anatomy and physiology. As was schematized in Fig. 1B, lower-order species like the mouse contain a kidney with a single renal papilla that is encased by the renal pelvic muscle. By contrast, as demonstrated in Fig. 6, higher-order mammals such as human and swine have kidneys with multiple renal papillae (see for example 6E, porcine UUT, papillae, pa). Moreover, the renal pelvis has branches of muscle that extend toward and encase each of these papillae. These units consisting of papillae encased by RPSM are termed calyces, thus giving rise to the designation multicalyceal UUT. Notably, in order to propel urinary waste away from the kidney, contractions of the multicalyceal UUT must propagate in an organized fashion from the proximal calyces distally through the RPSM and ureter ^32-34^. To date, however, the pacemaker mechanisms triggering multicalyceal UUT peristalsis remain elusive.

**Figure 6.**
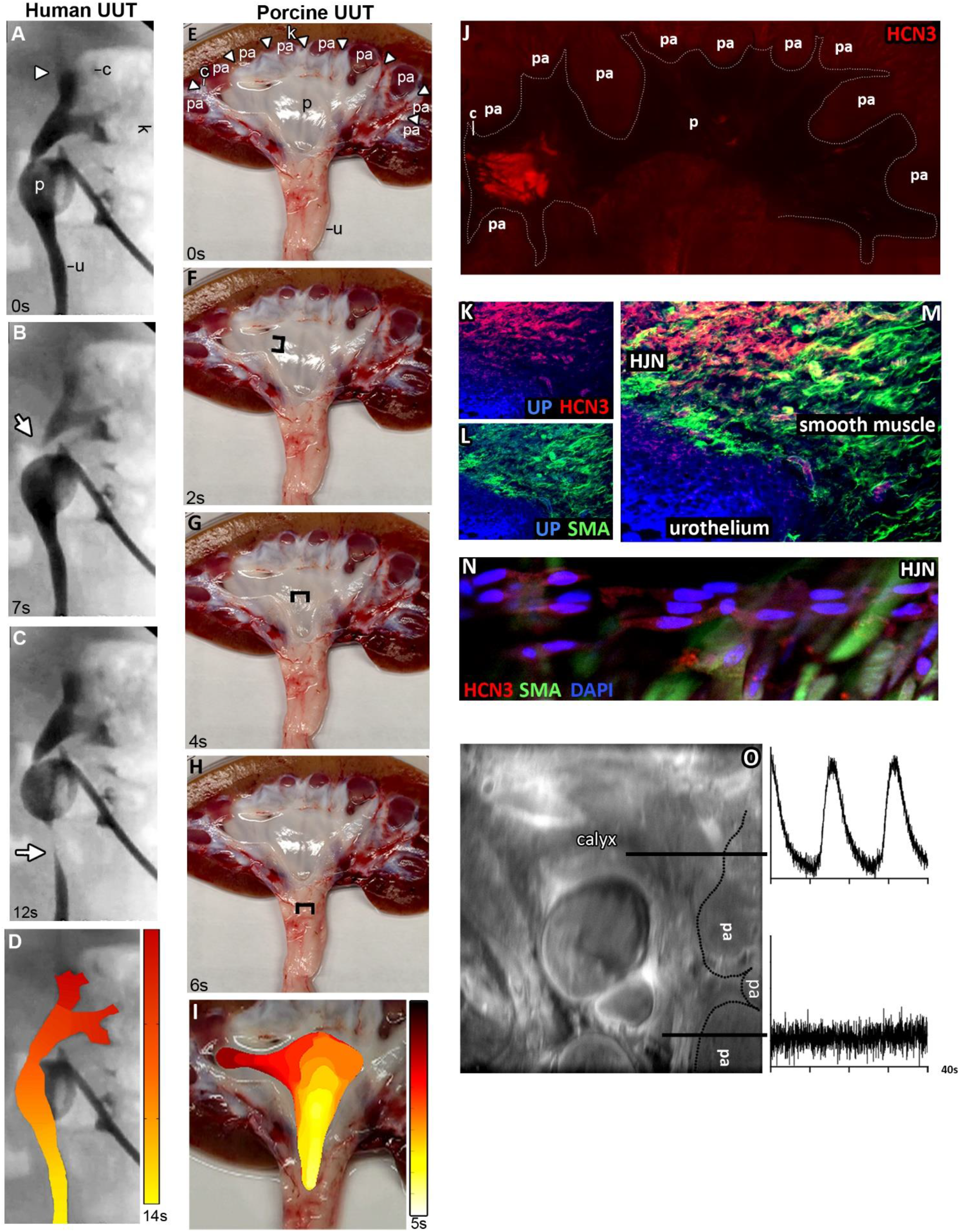
Calyces serve as the origin of peristalsis, elicit pacemaker depolarizations, and exhibit a conserved tissue architecture consisting of an HCN+ tissue layer. **A-D)** Representative still frames of live imaging of human multicalyceal UUT peristalsis via real-time fluoroscopic recordings. See supplemental movie 9 of live recording. Arrow head indicates origin of contraction at the calyx (A); arrows indicate point of contraction over time (B and C; D, isochronal map of contraction over time, from red initiation site to yellow). **E-I)** Representative still frames of live imaging of porcine UUT peristalsis. See supplemental movie 10 of live recording. Brackets indicate point of contraction over time (F-H; I, isochronal map of contraction over time, from red initiation site to yellow). **J)** Fluorimetric IF staining of 1mm thick porcine UUT sections labeled for HCN3 (K, red). **K-M)** Standard IF of calyx cryosections labeled for HCN3 (red, L and N), SMA (green, M and N) and uroplakin (blue, L-N). **N)** Confocal imaging of calyx tissue immunolabeled for HCN3 (red), SMA (green), and DAPI (blue). **O)** Detection of spontaneous pacemaker membrane depolarizations elicited at the calyx (top trace) via dual emission ratiometric optical mapping (bottom shows adjacent regions lacked pacemaker depolarizations). k, kidney; c, calyx; p, pelvis; u, ureter; pa, papilla; up, uroplakin; sma, smooth muscle actin.

To address the current gaps in knowledge of UUT myogenic physiology we assayed peristalsis of the human and porcine multicalyceal UUTs (Fig. 6). The human and porcine UUTs are homologous in their anatomy and physiology ^30, 31^, and the swine has been validated as key translational animal model system ^40, 41^. The rhythmic peristaltic contractile behavior of the human UUT was imaged via real-time fluoroscopic recording in patients undergoing nephrostograms where the contrast was applied and allowed to stabilize. Contractions emanated from the proximal calyx, and propagated in a coordinated fashion distally down the RPSM (p) to the ureter (u), as can be seen in representative time lapse images and supplemental Movie 9 (Fig. 6A-C, still images; D, isochronal map of contraction over time). These data provide a blueprint of multicalyceal UUT behavior, establishing that peristalsis persists as waves initiating at the calyces. Experimentation to better understand these observations in humans, however, was not feasible, and thus a translational animal model system was required to further investigate.

Whole kidneys and adjoined UUTs were isolated from swine, rinsed with Tyrode’s saline, and kidneys bisected along the sagittal plane within 5 min post-euthanasia. Bisection exposed the interior of the organ and enabled visualization of the intact UUT, including the calyx (c), renal pelvis (p) and ureter (u) segments (Fig. 6E). The preservation of the calyces was readily visualized, with each calyx exhibiting RPSM segments extending into the parenchyma of the kidney (6E, arrow heads). Explants were then placed in a bathing solution of Tyrode’s that washed over the tissue without submerging it, at 37°C and 5% CO_2_. As illustrated in supplemental Movie 10 and representative time lapse images (Fig. 6E-H; I, isochronal map of contraction over time), UUT explants prepared in this manner maintained their myogenic contractility, exhibiting spontaneous, rhythmic peristaltic contractions that propagate in a coordinated proximal-to-distal fashion from the calyx, down the renal pelvis and ureter. Thus, taken together, these live imaging data demonstrate that the calyx serves as the localized origin of peristalsis in both the porcine and human UUT.

Multicalyceal porcine UUTs were next assayed to define their expression of HCN pacemaker ion channels. One millimeter thick sagittal vibratome sections were cut to obtain an entire cross section of the UUT, and fluorimetric IF performed. We discovered HCN3 expression was conserved at the RPSM calyx of the UUT (Fig. 6J). To corroborate these findings, high magnification immunofluorescence of calyx cryosections was also performed (Fig. 6K-M). Calyces were stained for HCNs (red) in conjunction with calyceal tissue layer markers, namely the urothelium (Fig. 6K, uroplakin, blue,) that adjoins the renal papillae, and the UUT smooth muscle (Fig. 6L, SMA, green). These analyses confirmed the presence of a cluster of HCN+ cells, which formed a tissue layer integrated within the calyx RPSM (Fig. 6M, all merged). Indeed, IF and confocal microscopy of porcine UUTs enabled the visualization of HCN3+ cell clusters subadjacent to RPSM cells (Fig. 6N).

The electrical excitation of the multicalyceal UUT was assayed to define the pacemaker mechanisms driving UUT peristalsis. Dual emission ratiometric optical mapping of membrane potential was performed on proximal UUT tissues to visualize the generation of pacemaker potentials in real-time (Fig. 6O). UUTs were loaded with the membrane voltage sensitive dye Di-4-Anepps that exhibits depolarizing voltage-dependent shifts in the intensity of green (increased) and red (decreased) emission ^42, 43^. The shifts in the ratio of green/red emission of Di-4-ANEPPS follow action potential contours and delineate membrane potentials at the cellular and tissue levels ^44, 45^. To resolve pacemaker depolarizations from composite propagated electrical excitation, we inhibited excitation-contraction coupling with the L-type calcium channel blocker nifedipine. Inhibition of L-type calcium channels impedes smooth muscle excitation, and has been established to facilitate visualization of rhythmic pacemaker depolarizations ^46-48^. As can be seen in Fig. 6O, spontaneous rhythmic pacemaker depolarizations localized to the calyx of the UUT (top trace). By contrast, pacemaker depolarizations were not detectable in adjacent UUT tissues (bottom trace). Thus, pacemaker depolarizations are elicited at the calyx, the origin of contractions where peristalsis of the multicalyceal UUT initiates and where HCN channels are highly expressed.

We next determined if HCN channel activity is required for the generation of pacemaker depolarizations and coordinated peristalsis of the multicalyceal UUT (Fig. 7). The electrical and contractile activities of UUTs were assayed with and without HCN channel inhibition. Spontaneous, rhythmic pacemaker depolarizations were elicited at the calyx of control explants treated with vehicle (Fig. 7A; n = 7). Control explants also exhibited coordinated peristaltic contractions that initiated at the calyces and propagated distally down the UUT (Fig. 7B-G, brackets follow contractile site; supplemental Movie 11, live imaging of peristalsis; n= 5). By contrast, HCN channel block markedly perturbed the myogenic physiology of the UUT. Inhibition of UUTs with the ZD7288 abolished the generation of pacemaker depolarizations (Fig. 7H; n= 9). Consistent with a loss of pacemaker excitation, robust coordinated peristaltic contractions exhibited by UUTs were lost upon HCN channel block (supplemental Movie 12, before inhibition; supplemental Movie 13, post inhibition; n= 4). Instead, inhibited UUTs exhibited muscle twitching in localized segments (Fig. 7I-N, bracket marks site of twitch). These data demonstrate that HCN channels underlie the pacemaker activity that drives coordinated peristalsis of the multicalyceal UUT.

**Figure 7.**
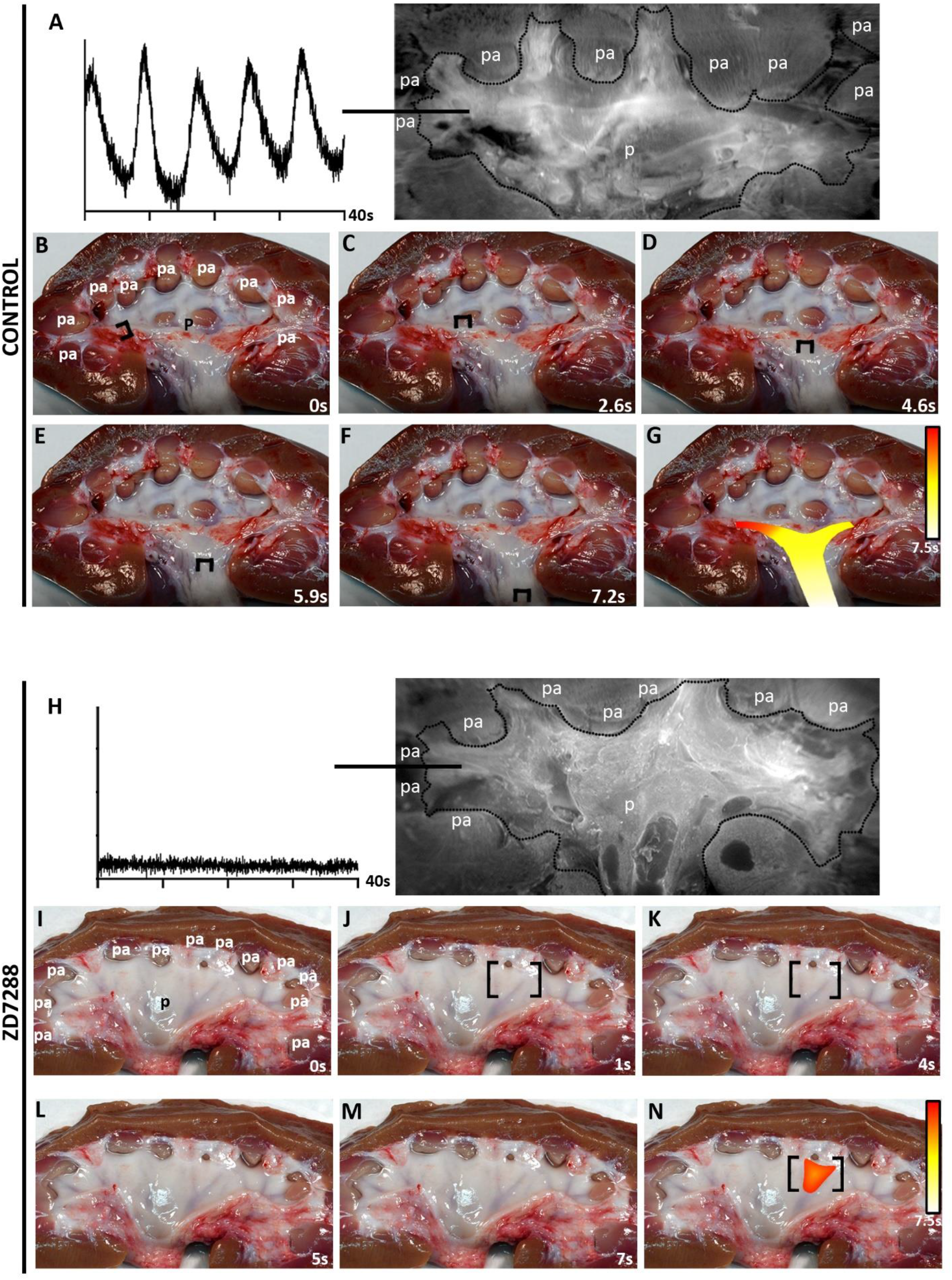
HCN channel conductance underlies pacemaker activity that triggers multicalyceal UUT peristalsis. **A-G)** Direct visualization of electrical and contractile excitation in control porcine UUTs. (A) Pacemaker depolarizations detected at the calyx of control porcine UUTs by dual emission ratiometric optical mapping. (B-G) Live imaging of coordinated, proximal to distal peristaltic contractions of control porcine UUTs (representative still images of supplemental movie 11, brackets mark site of contraction; G isochronal map of contraction over time, red initiation to yellow). **H-N)** Direct visualization of electrical and contractile excitation in porcine UUTs upon HCN channel block. (H) Pacemaker depolarizations were not detected in porcine UUTs inhibited with 100 µM ZD7288. (I-N) Live imaging of twitch like contractile activity in porcine UUTs inhibited with 100 µM ZD7288 (Supplemental movie 12, before inhibition; Supplemental movie 13, after inhibition; brackets mark the site of twitching; N, isochronal map of contraction over time, red initiation site to yellow). pa, papillae; p, renal pelvis.

Taken together, using multi-faceted imaging technologies, electrophysiological analyses, and both basic and translational animal models, we reveal a conserved HCN+ RPSM pacemaker mechanisms that drives peristalsis of the UUT in lower- and higher-order mammalian species.

## DISCUSSION

Morphological and functional analyses were used to reveal elements of a conduction system in RPSM. Imaging of the murine UUT revealed that the PKJ pacemaker is comprised of an HCN+ tissue layer integrated into the UUT muscular syncytium. HCN channel activity was required for rhythmic smooth muscle contractions driven by the PKJ. HCN+ cells expressed HCN channels on their plasmalemma, were selectable via immunopanning for HCNs, and exhibited the *I*_h_ pacemaker current characteristic of HCN channels. The latter addresses the previous lack of a detectable *I*_h_ current, which has been a central argument against HCN+ cells being UUT pacemakers ^19, 28^. Indeed, we demonstrate that 96% of cells purified from the PKJ by immunopanning exhibit the *I*_h_ pacemaker current. Moreover, an HCN+ tissue layer was conserved in calyx RPSM pacemaker tissue of the multicalyceal UUT. Though studies dating to the mid 1970’s – 1980’s described peristalsis initiating from the calyces, the molecular mechanisms, including the ion channels, underlying pacemaker activity of the calyces remained unknown ^32-34^. Utilizing our translational model system, we reveal that HCN channel conductance underlies calyx pacemaker depolarizations that trigger multicalyceal UUT peristalsis. Loss of HCN-dependent calyx pacemaker excitation resulted in a loss of coordinated peristalsis and resulted in muscle twitching. These data provide insight into studies showing HCN channel deficits in patients with aberrant UUT peristalsis. Although these studies envisioned a possible pacemaker role for HCN channels in human UUT disease, the authors noted a lack of human functional and electrophysiological evidence in their study and in the published literature precluded such a correlation ^27^. Taken together, these studies establish an evolutionarily conserved pacemaker mechanism in mammalian smooth muscle.

ICC cells have long been studied as pacemakers of the GI tract ^49-52^, and the recent identification of ICC-like cells expressing PDGFR-α in the UUT raised interest in this cell population. Indeed, based on their morphological characteristics, these cells have been proposed to be pacemakers ^28^. However, putative UUT pacemakers have been established to lack ANO1 channels ^26, 53^, whereas all ICC-like PDGFR-α+ cells express ANO1, thus raising questions regarding the identity of these cells. Indeed, those studies uncovered a disseminated distribution of PDGFR-α+ cells. For instance, in a subset of standard UUT cryosections these investigators found HCN+ cell clusters localized to the murine UUT pacemaker, consistent with our current findings and those of previously published studies of the UUT ^20-25^. By contrast, PDGFR-α+ cells were not restricted to UUT pacemaker tissues, but rather formed a cellular network that spanned the UUT, reminiscent of a vascular bed. Moreover, in these studies PDGFR-α+ cells were also shown to form an interconnected network surrounding the renal tubules, the site of the peritubular vasculature, further buttressing their possible role as accessory cells to the vasculature. We assayed for PDGFR-α+ cells in RPSM pacemaker tissues, distal UUT muscle, and kidney. We established that in all tissues PDGFR-α+ cells are mural cells associated with the vascular tree. Our findings in the kidney are consistent with well documented studies establishing PDGFR-α cells as renal vascular mural cells ^35^. Also consistent with the known phenotype of mural cells, PDGFR-α+ cells of the UUT associated with large caliber vessels, whereas subsets of smaller vessels lacked this mural cell coverage. Thus, our study resolves the identity of PDGFR-α+ cells of the UUT, establishing that they are vascular-associated mural cells.

Our studies revealed an evolutionarily conserved pacemaker mechanism that drives coordinated UUT peristalsis in mammals. It remains to be determined how these pacemaker tissues become specified, in particular in the multicalyceal UUT that exhibits a variable anatomy. For example, the number of calyces and their specific location in the UUT varies among individuals ^29^. Notably, hierarchical signaling by the T-box transcription factor family has been shown to play a critical role in cardiac pacemaker specification. By contrast, the mechanisms specifying the UUT pacemaker tissues remain elusive. Thus, analyses of the T-box gene family in the UUT may shed light on myogenic UUT physiology ^54^. Finally, it is well established that the cardiac conduction system consists not only of a primary SAN pacemaker, but also contains a secondary atrioventricular pacemaker ^1^. The presence of additional conduction system elements in the mammalian UUT remains a possibility, and it will be important to understand how these specialized units work together to permit coordinated peristalsis in the multicalyceal UUT.

In summary, we used basic and translational animal model systems to elucidate the presence of an evolutionarily conserved HCN+ pacemaker mechanism in mammalian smooth muscle. This provides evidence of functional conduction system elements that drive smooth muscle, akin to that of the SAN that drives contraction of the heart. Notably, HCN channelopathies are implicated in cardiac and neuronal pathologies, and may play a role in common UUT dysfunctions such as recurrent urinary tract infections and hydronephrosis. Thus, further translational studies into specialized conduction systems in the UUT and other smooth muscles will provide much needed insight into the behavior of this muscle type, and may pave the way for novel therapies to treat smooth muscle organ disorders.

## METHODS

### Animals

Adult mice were purchased from Taconic Farms (Germantown, NY, USA). Yorkshire and Micro Yucatan swine were purchased from Archer Farms (Darlington, MD, USA) and Sinclair Research (Auxvasse, MO, USA), respectively. All animals were housed at an AAALAC accredited facility at the Weill Medical College of Cornell University Animal Facility. All experiments were approved and carried out in accordance with the ethical guidelines and regulations stipulated by the Institutional Animal Care and Use Committee of Weill Medical College of Cornell University.

### Immunofluorescence

Immunofluorescence was performed based on our published studies ^23, 24^, with modification as follows: Samples were rinsed with PBS, and then permeabilized by treatment at 4°C with the following detergent solutions in sequence: 0.5% triton-x, 20% DMSO in PBS (TPD), followed by 0.5% triton-x, 0.1% Saponin, and 20% DMSO in PBS (TSPD). Time of permeabilizaton was dependent on tissue sample. Porcine UUTs were permeabilized in TPD for 1 week and TPSD for 1 week, whereas mouse UUTs were permeabilized in TPD at 4C for 72 hrs, followed by 72 hrs in TSPD at 4C. Tissues were then blocked in 10% normal serum in TSPD at 4°C. Blocking for porcine UUTs was 72 hrs, and mouse UUTs for 24 hrs. Samples were then incubated with antibodies in 0.5% triton-x, 1% DMSO in PBS. Antibodies used in this study were: anti-α-smooth muscle actin (clone 1A4, Sigma, at 1:2000), anti-endomucin (ab106100, Abcam, at 1:200), anti-Pdgfr-α (12-1401-81, ThermoFisher at 1:100), anti-HCN3 (APC-083, Alomone, at 1:500), and anti-uroplakin (H-180, Santa Cruz, at 1:50). Minimally cross-absorbed Alexa Flour secondary antibodies were used for primary antibody detections. Minimally cross absorbed AP-conjugated secondary antibody was used for fluorimetric IF, and signal detection done with Fast Blue (F3378, Sigma, .25mg/ml) and Naphtol-AS-MX-phosphate (N5000, Sigma, .25mg/ml) combination ^55^. Optical clearing was performed on whole tissues prior to confocal microscopy, based on ClearT2 ^56^. Tissues were first primed by treating with formamide, and then infused with increasing concentration of glycerol in combination with formamide. Confocal microscopy was done on a Zeiss LSM 800, and wide field fluorescence microscopy on a Zeiss Axio Observer or Nikon TE 200 mounted with CoolSnap-HQ and Hamamatsu CCD cameras, respectively. Image analyses were performed using MetaMorph (Molecular Devices) and Imaris Imaging Software (Bitplane).

### UUT explants

Murine UUT explants were prepared as previously reported ^20, 23^. In short, kidneys were isolated from 5 week old mice within 3 min post-euthanasia, and placed in Tyrode’s saline containing sodium bicarbonate and bisected alongside the renal papillae to expose the intact UUT. Explants were then transferred to 24-mm Costar Transwell plates (Corning) containing Tyrode’s and placed on an inverting rocker within a 5% CO2 incubator, so that Tyrode’s bathes over the explants without completely submerging them. For microdissection of whole proximal UUTs and UUT segments, kidney parenchyma surrounding the UUT was dissected away using a tungsten wire tool and spring scissors. For Porcine UUTs, specific pathogen free swine (*Sus scrofa*) 8-10 weeks old were sedated with tiletamine (10 mg/ml) and zolazepam (10 mg/ml) combination given intramuscularly into the right quadriceps at a rate of 1 ml/10 pounds followed by administration via auricular vein of pentobarbital (390 mg/ml) and phenytoin (50mg/ml) combination at a dose of 1 ml/10 pounds. Once cardiorespiratory arrest was confirmed via auscultation, the ventral abdominal wall was incised and both kidneys with adjoined ureters were harvested within a 5 min period. Isolated tissues were rinsed with Tyrode’s, and kidneys bisected along the frontal plane, anterior to the renal pelvis to expose the intact UUT. Explants were then placed in custom containers housed within a 5% CO_2_ incubator, on an inverting rocker to enable the explant to be washed over with Tyrode’s but not submerged. Nifedipine (Sigma) stock solution of 100 mM in DMSO was brought to a final concentration of 10 µM to inactivate UUT smooth muscle, as has been previously described ^23, 46, 47^. ZD7288 (Tocris Bioscience) stock solution of 100 mM was made in PBS and stored at - 20°C. For HCN channel inhibition of explants, ZD7288 was diluted 1:2 with Hybri-max DMSO, and then brought to a final concentration of 100 µM in Tyrode’s. Fresh solution of ZD7288 was added every 30 min for a total of 2hrs. Real time videomicroscopy of UUT peristalsis was done using a Nikon D810A camera.

### Super-resolution STORM

STORM microscopy provides sub-diffraction limit resolution by repeated imaging of stochastically activating fluorescent probes, with super-resolution images of single molecules reconstructed from the calculated localization of individual fluorophores ^57, 58^. PKJ pacemaker tissue sections were labeled with anti-HCN3 antibody, followed by Alexa 647 conjugated secondary antibody. STORM analysis of Alexa 647 was performed by stochastically imaging spontaneous fluorescence using a 70mW Agilent 647nm laser line. Experiments were performed on a Nikon Eclipse Ti microscope with a Nikon Plan Apo TIRF 100×/1.45 N.A. objective, an Andor iXon EMCCD camera, and an NSTORM Quad cube (Chroma). Samples were imaged in a buffer containing 50 mM Tris, pH 8, 10 mM NaCl, 10% (wt/vol) glucose, 500 μg/mL glucose oxidase, 10 mM MEA, and 4 mg/mL catalase. The laser angles were set for total internal fluorescence to restrict visualization to the coverslip proximal portion of the sample. Thirty-five thousand images were collected at an average frame rate of 56 frames/second and fitted with 2D Gaussian function and drift corrected using the Nikon NIS Elements software.

### Primary Cell Isolation

Kidneys with adjoined ureters were isolated from mice and placed in Leibovitz’s L-15 medium (Corning). Kidneys were bisected by cutting longitudinally alongside the renal papillae to expose the center of the organ and the UUT. PKJ tissues were micro-dissected out and rinsed in fresh L-15 media. Isolated PKJ tissues were then enzymatically digested with collagenase type IV (1mg/ml, Stemcell Technologies) for one hour at 37^0^C. Following digestion, PKJ tissues were gently dispersed by manual pipetting to obtain single cells. Cell suspensions were added to 10 ml of advanced DMEM (ThermoFisher) supplemented with 1% GlutaMax (DMEM-G), and centrifuged at 300 x g for 10 minutes at room-temperature. The supernatant was decanted, cells re-suspended in DMEM-G, and resultant primary cells directly used in patch-clamp and immunopanning experiments.

### Immunopanning

Immunopanning protocols were developed based on the work of Barres ^59^. Petri-dishes were coated with antibodies at a concentration of 6µg/ml in Tris-HCl, pH 9.5, overnight at 4°C. For depletion of immune cells by their FC receptors, we prepared negative selection plates with minimally absorbed, affinity purified anti-rabbit antibody (Jackson Immunoresearch). For selection of HCN3^+^ cells, we prepared plates with anti-HCN3 antibody (extracellular anti-HCN3, Alomone, APC-083). Primary cells were incubated on negative selection plates for 1 hour at room temperature, with manual rocking of plates done every 15 min. Supernatant with unbound cells was gently transferred to selection plates by decanting. Decanted cells were incubated on selection plates for 1 hour at room temperature, with manual rocking done every 15 min. Unbound cells were gently aspirated off of selection plates, and selection plates washed with DMEM 2x times at room temperature. Selected, bound cells were then removed from plates by manual pipetting, and then centrifuged at 300 x g for 10 minutes. Cells were re-suspended and incubated in DMEM-G, with 1% fetal bovine serum (ThermoFisher), at 37 °C, 5% CO_2_ for 1 hr., and then used in patch clamp studies.

### Electrophysiology

Primary cells were analyzed using the whole-cell patch-clamp technique. Pipettes were prepared from hematocrit capillary glass (VWR Scientific) using a three-stage vertical puller, and had resistances of 2-10 MΩ. The standard extracellular solution was composed of (in mM):140 NaCl, 5 KCl, 0.33 NaH_2_PO_4_, 0.5 MgCl_2_, 1.8 CaCl_2_, 5 HEPES, 5.5 glucose, 0.5 BaCl_2_, pH=7.4, adjusted with NaOH. The pipette solution contained (in mM): 100 aspartic acid, 5 Na_2_ATP, 0.1 Mg GTP, 5 EGTA, 5 HEPES, 20 KCl, 1 MgCl_2_, pH 7.2, adjusted with KOH. The final K concentration in the pipette solution was 120 mM. The extracellular solution superfusing the cells was controlled by a rapid exchange system (AutoMate Scientific). ZD7288 was purchased from Tocris Biosciences (MN, USA). The experiments were carried out at 33-35 °C. After establishment of a whole-cell clamp, the stability of currents was assessed for at least two minutes. Recordings in which the I_h_ showed significant rundown were discarded. Data were acquired at 10 kHz using an EPC-7 patch-clamp amplifier (HEKA) and digitized with a Digidata 1332A interface (Molecular Devices). Data were filtered at 0.5 kHz and analyzed with pClamp10 (Molecular Devices) and Origin 6 software. For determination of the voltage of half-maximal activation (V_0.5_) currents were elicited by hyperpolarizing the membrane for 5-10s to voltages from - 150 mV to -20 mV (in 10 mV increments) followed by a 500 ms step to -130 mV or -150 mV. Initial current amplitude, determined immediately after the decay of the capacitance transient, were normalized to the value at -130 mV and plotted as a function of the preceding membrane potential. The data points are fitted with the Boltzmann function (equation 1),

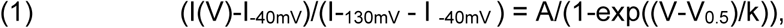

I_-40mV_ is an offset caused by a non-zero holding current, V is the test potential, V_0.5_ is the voltage of half-maximal activation, A is the maximal value, and k is the slope factor. The *I*_h_ amplitude was measured as a difference between the I_ins_ (instantaneous current) and I_ss_ (steady-state current), and data are expressed as mean ± SEM.

### Optical Mapping

Optical mapping of electrical excitation was performed as previously described ^20, 23, 60^. In short, UUT explants were loaded with the membrane voltage sensitive dye di-4-ANEPPS (Invitrogen) that exhibits depolarizing voltage dependent shifts in green emission (increased intensity) and red emission (decreased intensity) wavelengths ^42, 43^. Dual-wavelength changes of green/red emission ratios follow action potential contours at cellular and tissue levels ^44, 45^. Murine UUT explants were loaded with di-4-ANEPPS (20 µM) for 10 min within a 5% CO_2_ incubator. Porcine UUT explants were loaded with di-4-ANEPPS (20 µM) for 15 min within a 5% CO_2_ incubator. Fluorescent intensity of Di-4-ANEPPS was monitored by excitation at 470±20 nm (OptoLED light source, Cairn Research) and detection of separated (OptoSplit II, Cairn Research) bandpass-filtered green emission (502-557 nm, Chroma Technology) and red emission (642-708 nm, Chroma Technology) using a NEO-CMOS camera (Andor Technology) mounted on a MVX10 macroscope (Olympus) at an acquisition speed of 50 Hz. Data was acquired and analyzed by Solis (Andor) and MetaFluor (Molecular Devices) imaging software, respectively.

## ACKNOWLEDGEMENTS

We thank Dr. Lauretta Lacko for thoughtful suggestions on the manuscript, Dr. Bob Switzer for technical assistance processing porcine tissues, and Mr. David Gyorgi for advice on image analyses. This work was supported by the NIH NIDDK R21 DK116171 to R.H., and R01 DK 111380 to L.P.

## Data Availability

All data needed to evaluate the conclusions in the paper are present in the paper and/or the Supplementary Materials

## AUTHOR CONTRIBUTIONS

Study conception, R.H; Experiment Design, R.H., L.P., L.Y., C.S., R.R; Experiments, R.H., L.Y., K.B., C.S., R.R.; Intellectual Contributions, R.H., L.P, T.E., R.R.; Manuscript Writing, R.H. with input from L.P., T.E., R.R.

## DECLARATION OF INTERESTS

None

**Supplemental Movie 1**. Imaging of the intact murine atria and sinoatrial node

**Supplemental Movie 2**. The myogenic peristalsis of the murine UUT

**Supplemental Movie 3**. Imaging of intact PKJ pacemaker tissues

**Supplemental Movie 4**. The isolated proximal UUT continues to exhibit myogenic peristalsis

**Supplemental Movie 5**. Dissected PKJ tissues exhibit peristalsis

**Supplemental Movie 6**. UUT segments dissected away from the PKJ lack myogenic peristalsis

**Supplemental Movie 7**. Myogenic PKJ contractility prior to inhibition

**Supplemental Movie 8**. HCN channel block abolishes myogenic PKJ contractility

**Supplemental Movie 9**. Live imaging of multicalyceal human UUT peristalsis

**Supplemental Movie 10**. Live imaging of multicalyceal porcine UUT peristalsis

**Supplemental Movie 11**. Control porcine UUTs exhibit rhythmic myogenic peristalsis

**Supplemental Movie 12**. Porcine UUTs exhibit rhythmic myogenic peristalsis prior to HCN channel block

**Supplemental Movie 13**. HCN channel block abolishes myogenic peristalsis exhibited by the porcine UUT

---

**Supplemental Figure 1.**
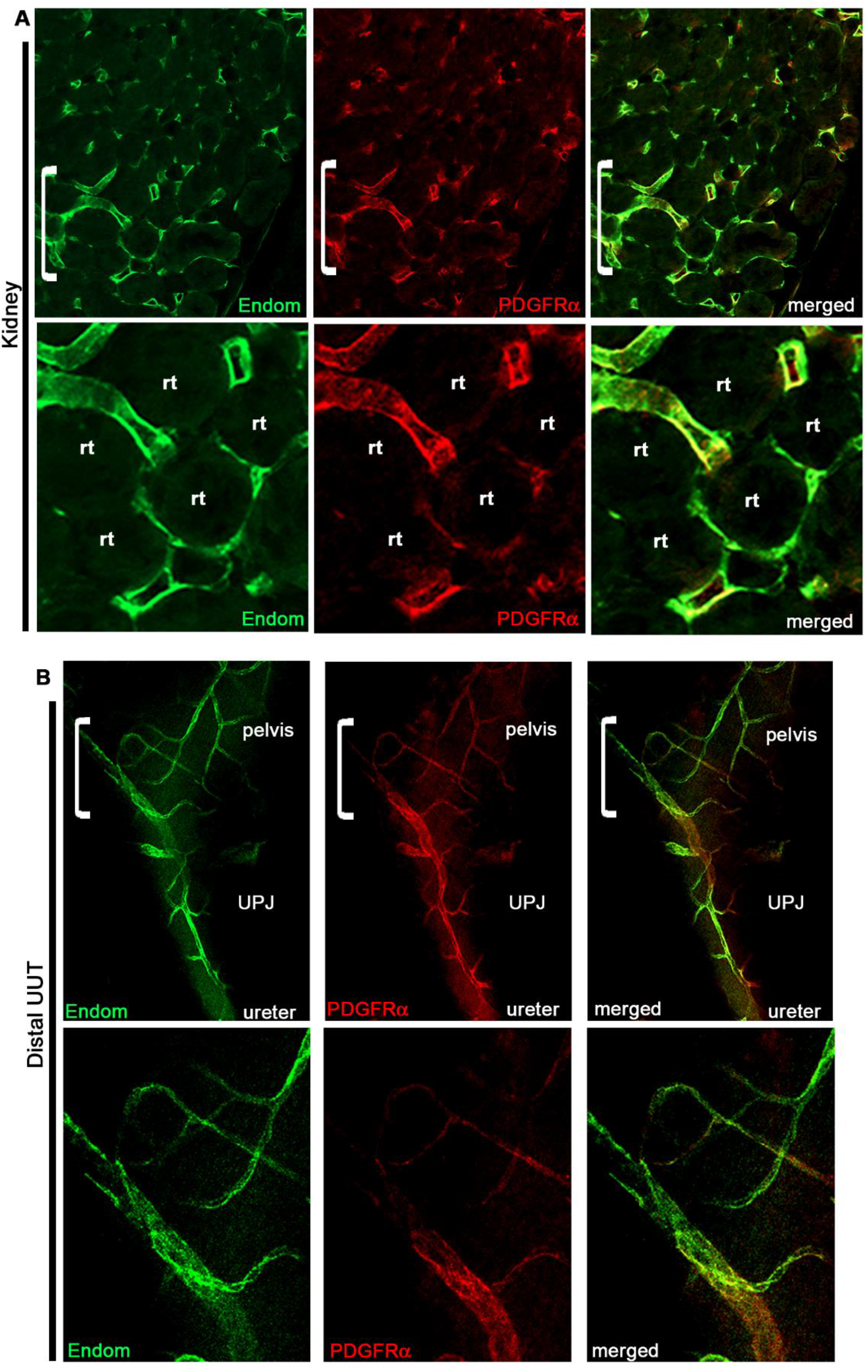
PDGFR-α cells of the kidney and distal UUT are vascular associated mural cells. **A, B) T**he cell identity of PDGFR-α+ cells in the kidney (A) and distal UUT (B). Tissues were immunostained for the vasculature (vascular endothelium, endomucin, green) and PDGFR-α (red). PDGFR-α cells were established to be vascular associate mural cells that overlay the vasculature in both the kidney and distal UUT. Brackets in A and B mark regions assayed at higher magnification (bottom panels of A and B).

